# Task complexity interacts with state-space uncertainty in the arbitration process between model-based and model-free reinforcement-learning at both behavioral and neural levels

**DOI:** 10.1101/393983

**Authors:** Dongjae Kim, Geon Yeong Park, John P. O’Doherty, Sang Wan Lee

## Abstract

A major open question concerns how the brain governs the allocation of control between two distinct strategies for learning from reinforcement: model-based and model-free reinforcement learning. While there is evidence to suggest that the reliability of the predictions of the two systems is a key variable responsible for the arbitration process, another key variable has remained relatively unexplored: the role of task complexity. By using a combination of novel task design, computational modeling, and model-based fMRI analysis, we examined the role of task complexity alongside state-space uncertainty in the arbitration process between model-based and model-free RL. We found evidence to suggest that task complexity plays a role in influencing the arbitration process alongside state-space uncertainty. Participants tended to increase model-based RL control in response to increasing task complexity. However, they resorted to model-free RL when both uncertainty and task complexity were high, suggesting that these two variables interact during the arbitration process. Computational fMRI revealed that task complexity interacts with neural representations of the reliability of the two systems in the inferior prefrontal cortex bilaterally. These findings provide insight into how the inferior prefrontal cortex negotiates the trade-off between model-based and model-free RL in the presence of uncertainty and complexity, and more generally, illustrates how the brain resolves uncertainty and complexity in dynamically changing environments.

**SUMMARY OF FINDINGS:** - Elucidated the role of state-space uncertainty and complexity in model-based and model-free RL.

- Found behavioral and neural evidence for complexity-sensitive prefrontal arbitration.

- High task complexity induces explorative model-based RL.

## INTRODUCTION

An influential computational account of reward-related learning and decision-making built on the basis of a large body of empirical evidence suggests that there exists two distinct mechanisms for controlling instrumental actions: a model-free RL system that learns values for actions based on the history of rewards obtained on those actions (Balleine and Dickinson, 1998; Dickinson, 1985; Graybiel, 2008), and a model-based RL (MB) system that computes action-values flexibly based on its knowledge about state-action-state transitions incorporated into an internal model of the structure of the world (Doya et al., 2002; Kuvayev et al., 1996). These two systems have different relative advantages and disadvantages for a behaving agent. For example, while model-free RL can be computationally cheap and efficient, it achieves this at the cost of a lack of flexibility, thereby potentially exposing the agent to inaccurate behavior following a change in the subjective value of the goal or outcome, or a rapid change in environmental contingencies (Balleine and Dickinson, 1998; Daw et al., 2005; Dickinson, 1985). On the other hand, model-based RL is computationally expensive as it requires active computation of expected values and planning, but retains flexibility in that it can rapidly adjust a behavioral policy following a change in the features of the outcome or environmental contingencies. The theoretical trade-offs that exist between these two systems has, alongside empirical evidence for the parallel existence of these two modes of computation in the behavior of animals and humans (Akam et al., 2015; Balleine and Dickinson, 1998; Balleine and O’Doherty, 2010; Beierholm et al., 2011; Daw et al., 2011; Doll et al., 2012, 2016; Gläscher et al., 2010; Gremel and Costa, 2013; Gruner et al., 2016; Hasz and Redish, 2018; Kool et al., 2017; Linnebank et al., 2018; McDannald et al., 2011; Miller et al., 2017; Russek et al., 2017; Sang Wan Lee et al., 2013; Simon and Daw, 2011; Skatova et al., 2013; van Steenbergen et al., 2017; Wunderlich et al., 2012; Yin and Knowlton, 2004; Yin et al., 2005), prompted interest in elucidating how it is the trade-off between these systems is actually managed in the brain.

One influential proposal is that there exists an arbitration process that allocates control to the two systems according to various criteria (Daw et al., 2005; Kool et al., 2017; Pezzulo et al., 2018). One variant of this theory suggests that estimates about the uncertainty in the predictions of the two systems mediates the trade-off between the respective controllers, such that under situations where the model-free system has unreliable predictions, the model-based system is assigned greater weight over behavior, while under situations where the model-free system has more accurate predictions, it will be assigned greater behavioral control (Daw et al., 2005; Lee et al., 2014).

One of the challenges in building a computationally and biologically plausible theory of the arbitration process, is the necessity that the arbitration process should not by itself be so computationally expensive as to render the relative savings in computational cost associated with being model-free vs between model-based to be rendered moot. For instance, for uncertainty-based arbitration, the computation of uncertainty might involve a computationally expensive process, while accurately estimating the potential gain to being model-based could under some implementations, actually require model-based computations to estimate that potential gain, which in many situations would defeat the purpose of the trade-off i.e. it could be more efficient to just remain model-based and dispense with arbitration altogether. To this end, practical computational theories of arbitration have examined computationally cheap approximations that might be utilized to mediate the arbitration process. According to Lee et al., uncertainty is approximated via a mechanism that tracks cumulative predictions errors induced in the two systems (Lee et al., 2014). Model-free uncertainty is approximated via the average accumulation of reward-prediction errors, while model-based uncertainty is approximated via the accumulation of errors in state-prediction (so called state-prediction errors).

Utilizing this framework, Lee et al., examined the neural correlates of the arbitration process (Lee et al., 2014). In that study, a region of bilateral ventrolateral prefrontal cortex was found to track the reliabilities (an approximation of the inverse of uncertainty) in the predictions of the two systems, consistent with a role for this brain region in the arbitration process itself. However, as alluded to above, the relative uncertainty or reliability in the predictions of the two systems is only one-component of the trade-off between the two controllers. Another equally important element of this trade-off is the relative computational cost of engaging in model-based control.

The goal of the present study is to investigate the role of computational cost in the arbitration process, alongside relative uncertainty. We experimentally manipulated computational cost by means of adjusting the complexity of the planning problem faced by the model-based controller. This was achieved by subjecting participants to a multi-step Markov Decision Problem in which the number of actions available in each state was experimentally manipulated. In one condition, which we called low complexity, two actions were available, while in another condition, which we called high complexity, four actions were available. In addition to manipulating complexity, we also, as in our previous study (Lee et al., 2014), manipulated the uncertainty in the state-space, by utilizing a state-transition structure in the MDP that invoked high levels of state-prediction errors (i.e. one where the transitions are maximally uncertainty), and a transition structure where the transitions are either high or low in uncertainty. Thus, we manipulated two variables in a factorial design: state-space complexity (low vs high), and state-space uncertainty (low vs high).

This design allowed us to investigate the ways in which uncertainty and complexity interact to drive the arbitration process, thereby allowing us to assess the interaction between (model-based) reliability and at least one simple proxy of computational cost. While participants were undergoing this novel behavioral task, we also simultaneously measured brain activity with fMRI. This allowed us to investigate the contribution of state-space complexity alongside state-space uncertainty in mediating arbitration at both behavioral and neural levels. In order to accommodate the effects of task complexity in the arbitration process, we extended our previous arbitration model to endow this arbitration scheme with the capability of adjusting the arbitration process as a function of complexity. On the neural level, we focused on the ventrolateral prefrontal cortex as our main brain region of interest, given this was the main region implicated in arbitration in our previous study. We hypothesized that an arbitration model which is sensitive to both the complexity of the state-space, and the degree of uncertainty in the state-space transitions would provide a better account of behavioral and fMRI data than would an arbitration model that was sensitive only to state-space uncertainty. We further hypothesized that under conditions of high state-space complexity, the model-based controller would be selected against, because the complexity of the state-space would overwhelm the planning requirements of the model-based system, forcing participants to rely instead on model-free control. Our findings support our first hypothesis, and partially support our second hypothesis.

## RESULTS

### Markov decision task with varying degrees of uncertainty and complexity

To investigate the role of uncertainty and complexity in arbitration control, we designed a novel two-stage Markov decision process task (MDP; Figure 1A), in which we systematically manipulated two task variables across blocks of trials, state-transition uncertainty, and state-space complexity (Figure 1B). The amount of state-transition uncertainty is controlled by means of the state-action-state transition probability. The state-action-state transition probability varies between the two conditions: high uncertainty (0.5 versus 0.5) and low uncertainty (0.9 versus 0.1). Switching between the two uncertainty conditions is designed to effect a change in the average amount of state prediction errors (SPE), thereby effectively translating into the reliability of the MB system. For instance, the high uncertainty condition will elicit a large amount of SPE, essentially resulting in a decrement in MB prediction performance. In the low uncertainty condition, the SPE will decrease or stay low on average as the MB refines an estimate of the state-action-state transition probabilities. On the other hand, the performance of MF is less affected by the amount of state-transition uncertainty (Daw et al., 2005). The second variable, the number of available choices, is intended to manipulate the task complexity. The total number of available choices is two and four in the low and high complexity condition, respectively. To prevent state-space representations from being too complex, we limit the number of available choices to two in the first stage of each trial, while the choice availability in the 2^nd^ stages are either set to two or four. The manipulation of choice availability creates wide variability in the number of ways to achieve each goals, causing the difficulty level on each trial to range from easy to arduous. This design therefore provides four different types of conditions (*low/high x uncertainty/complexity*; see Figure S1). Participants then make sequential choices in order to obtain different colored tokens.

**Figure 1.**
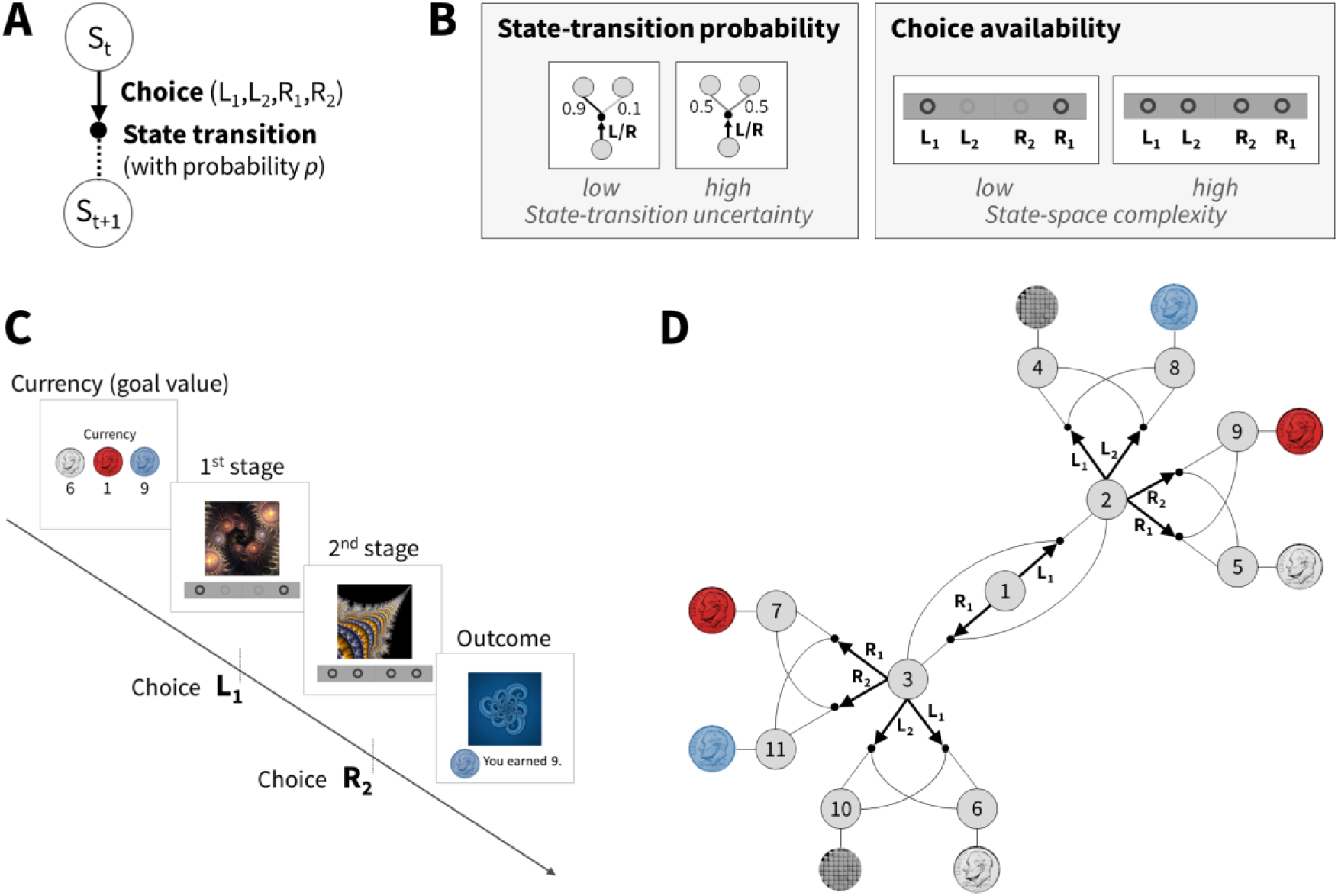
Task Design. **(A)** Two-stage Markov decision task. Participants choose between two to four options, followed by a transition according to a certain state-transition probability *p*, resulting in participants moving from one state to the other. The probability of a successful transitions to a desired state is proportional to the estimation accuracy of the state-transition probability, and it is constrained by the entropy of the true probability distribution of the state-transition. For example, the probability of a successful transition to a desired state cannot exceed 0.5 if *p*=(0.5, 0.5) (the highest entropy case). **(B)** Illustration of experimental conditions. (Left box) A low and high state-transition uncertainty condition corresponds to the state-transition probability p=(0.9,0.1) and (0.5,0.5), respectively. (Right box) The low and high state-space complexity condition corresponds to the case where two and four choices are available, respectively. In the first state, only two choices are always available, in the following state, two or four options are available depending on the complexity condition. **(C)** Participants make two sequential choices in order to obtain different colored tokens (silver, blue, and red) whose values change over trials. On each trial, participants are informed of the “currency”, i.e. the current values of each token. In each of the subsequent two states (represented by fractal images), they make a choice by pressing one of available buttons (L1, L2, R1, R2). Choice availability information is shown at the bottom of the screen; bold and light grey circles indicate available and unavailable choices, respectively. **(D)** Illustration of the task. Each grey circle indicates a state. Bold arrows and lines indicate participants’ choices and subsequent state-transition according to the state-transition probability, respectively. Each outcome state (state 4-11) is associated with a reward (colored tokens or no token represented by a grey mosaic image). The reward probability is 0.8.

Another feature of the MDP is that on each trial participants could take actions in order to obtain one of three different tokens, a silver, red or blue token (Figure 1C). On each trial, the relative value of the tokens, in terms of the rate of exchange of each token for real world money (US cents), was flexibly assigned, as revealed at the beginning of each trial. So for example, on a given trial, the silver token if won on that trial might yield 1 US cents, the red token, 9 US cents, and the blue token 3 cents, while the allocations could be different on the next trial. This design feature is intended to induce trial-by-trial changes in goal-values, thereby also inducing variance in reward-prediction errors and hence the reliability of MF across trials.

Twenty-four adult participants (twelve females, age between 19 and 55) performed the task, and among them 22 participants were scanned with fMRI. The task performance of subjects in terms of both the total amount of earned reward and the proportion of optimal choices is significantly greater than chance level in all conditions (t-test; p<1e-5).

### Computational model of arbitration control incorporating uncertainty and complexity

In our computational model, the dynamical interaction between MB and MF RL is based on a two-state transition model (Dayan and Abbott, 2001) (see Figure 2). Note that this type of recurrent structure has been previously found to account well for the arbitration process between MB and MF RL in both behavioral (Wang et al., 2018; Lee et al., 2014) and fMRI data (Lee et al., 2014). In this model, each state depends on its previous state (an endogenous input) and inputs from the environment (exogenous variables), such as a state, reward, and perceived task complexity. In this model, preference for MB and MF RL, indicated by the model choice probability (P_MB_), is a function of prediction uncertainty and task complexity. The prediction uncertainty refers to estimation uncertainty about state-action-state transitions and rewards. They are computed based on the state prediction error (SPE) and the reward prediction error (RPE), the key variables for MB and MF learning, respectively. We specifically hypothesized that task complexity influences the transition between MB and MF. Note that when the environment is perfectly stable (i.e., a fixed amount of state-transition uncertainty and a fixed level of task complexity), the particulars of this model converge to a stable mixture of MB and MF RL (Daw et al., 2011).

**Figure 2.**
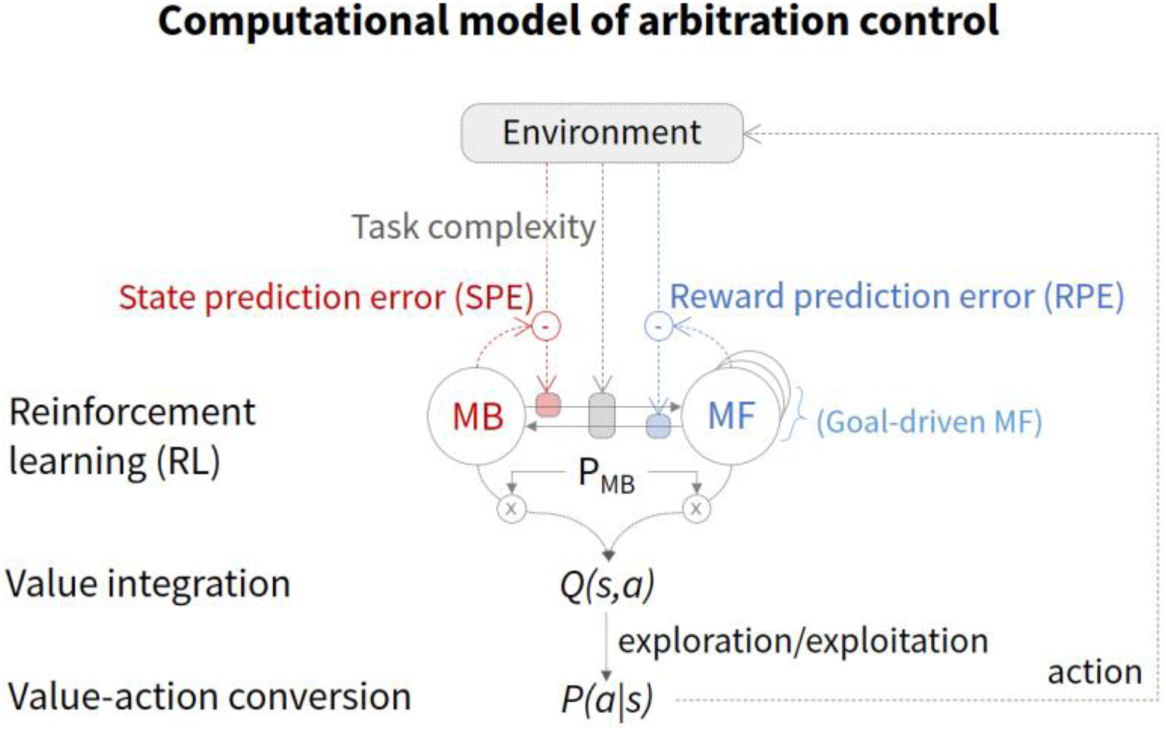
Computational model of arbitration control incorporating uncertainty and complexity. The circle-and-arrow illustration depicts a two-state dynamic transition model, in which the current state depends on the previous state (an endogenous variable) and input from the environment (exogenous variables). The environmental input includes the state-transition which elicits state prediction errors (SPEs), rewards that elicit reward prediction error (RPEs), and the task complexity. The arrow refers to the transition rate from MB to MF RL or vice versa, which is a function of SPE, RPE, and task complexity. The circle refers to the state, defined as the probability of choosing MB RL (P_MB_). Q(s,a) refers to the values of the currently available actions (a) in the current state (s). The value is then translated into action, indicated by the action choice probability P(a|s).

The process of our computational model is described as follows: first, in response to the agent’s action on each trial, the environment provides the model with the state-action-state transition, token values, and task complexity. These observations are then used to compute the transition rates (MB⟶MF and MF⟶MB), which subsequently determines the model choice probability PMB. Second, the model integrates MB and MF value estimations to compute an overall integrated action value (Q(s,a) of Figure 2), which is subsequently translated into an action (P(a|s) of Figure 2). It is noted that we use this framework to formally implement various hypotheses about the effect of uncertainty and complexity on RL. For instance, the configuration of the model that best accounts for subjects’ choice behavior would specify the way people combine MB and MF RL to tailor their behavior to account for the degree of uncertainty and complexity of the environment.

### Effect of uncertainty and complexity on reinforcement learning

To determine how task complexity is embedded into the arbitration control process, we formulated a variety of possible model implementations, which we could then fully permute and test in a large scale model comparison, described as follows:

The first factor incorporated into the hypothesis set is the effect of complexity on arbitration control. We considered a number of different ways in which complexity could impact on the allocation of control between MF and MB systems. The first of these variables was the type of modulation, i.e. whether modulation was excitatory or inhibitory. That is, does complexity influence on the arbitration process positively or negatively? (see Methods – subsection (1) Sign of modulation). The second variable was the direction of modulation, that is does complexity effect the transition of behavioral control from model-free to model-based, from model-based to model-free, or does it affect the transition in both directions (see Methods – subsection (2) Direction of modulation). We then also considered the extent to which the effects of uncertainty also interacted with the effects of complexity, specifying three possibilities: the version that there is no interaction between complexity and uncertainty, and the other two versions that these two variables interact in different ways (see Methods – subsection (3) Type of modulation).

The second factor is the effect of complexity on the implementation of the choice itself. That is, we tested whether task complexity impacted on the soft-max choice temperature that sets the stochasticity of the choices of the integrated model. For this we compared the case in which there is no effect of complexity on the choice temperature parameter (Null), and a case in which increasing complexity increases the degree of explorative choices, and when increasing complexity decreases the degree of explorative choices (see Methods – subsection 3. Effect of complexity on exploration).

Another factor that we considered concerns the implementation of the model-free RL algorithm. Recall that on the beginning of each trial, the participant is presented with the current value of each of the three tokens. One possible implementation of model-free RL is that it could ignore those token values completely, and treat each trial the same irrespective of what values are signaled to accrue to the particular tokens. However, given these tokens are salient stimuli that signal specific states in their own right, such a possibility seems unlikely. Rather it seems more plausible that the token values would be embedded into the state space itself, on which the model-free RL agent learns. Accordingly, we built a modified model-free RL algorithm that divided up the task into three unique state-space representations depending on which token was the dominant goal (which depends on which token was the most valuable on a given trial), a model variant we call the 3Q model. Yet another, albeit much less plausible possibility is that 3 completely separate model-free strategies exist to learn about the separate values of each possible goal, which we call the 3MF model. Thus, we tested 3 classes of model-free agent, ranging from a simple model-free agent that doesn’t differentiate between token states, a model that treats the most valuable token as identifying one of three relevant subsets of the state-space (red token goal, blue token goal and silver token goal) and a model that assumes 3 independent model-free agents for each selected goal.

The effect of uncertainty on the transition rate was implemented based on our previous model and findings (Lee et al., 2014). Each of these factors (and sub-factors) was fully tested in each possible combination with each other factor, rendering a total of 117 model variants, that also included as a baseline model, the original arbitration scheme from our 2014 paper that only incorporated uncertainty as a variable and not complexity (see Figure 2). We then compared the extent to which each of those models could explain the behavioral data using across all of these models. For this, we fit each version of the model to each individual subject’s data, and then ran a Bayesian model selection (Stephan et al., 2009), with an exceedance probability *p*>0.99 (Figure 3; For more details, see Model Comparison in Supplemental Experimental Procedures).

**Figure 3.**
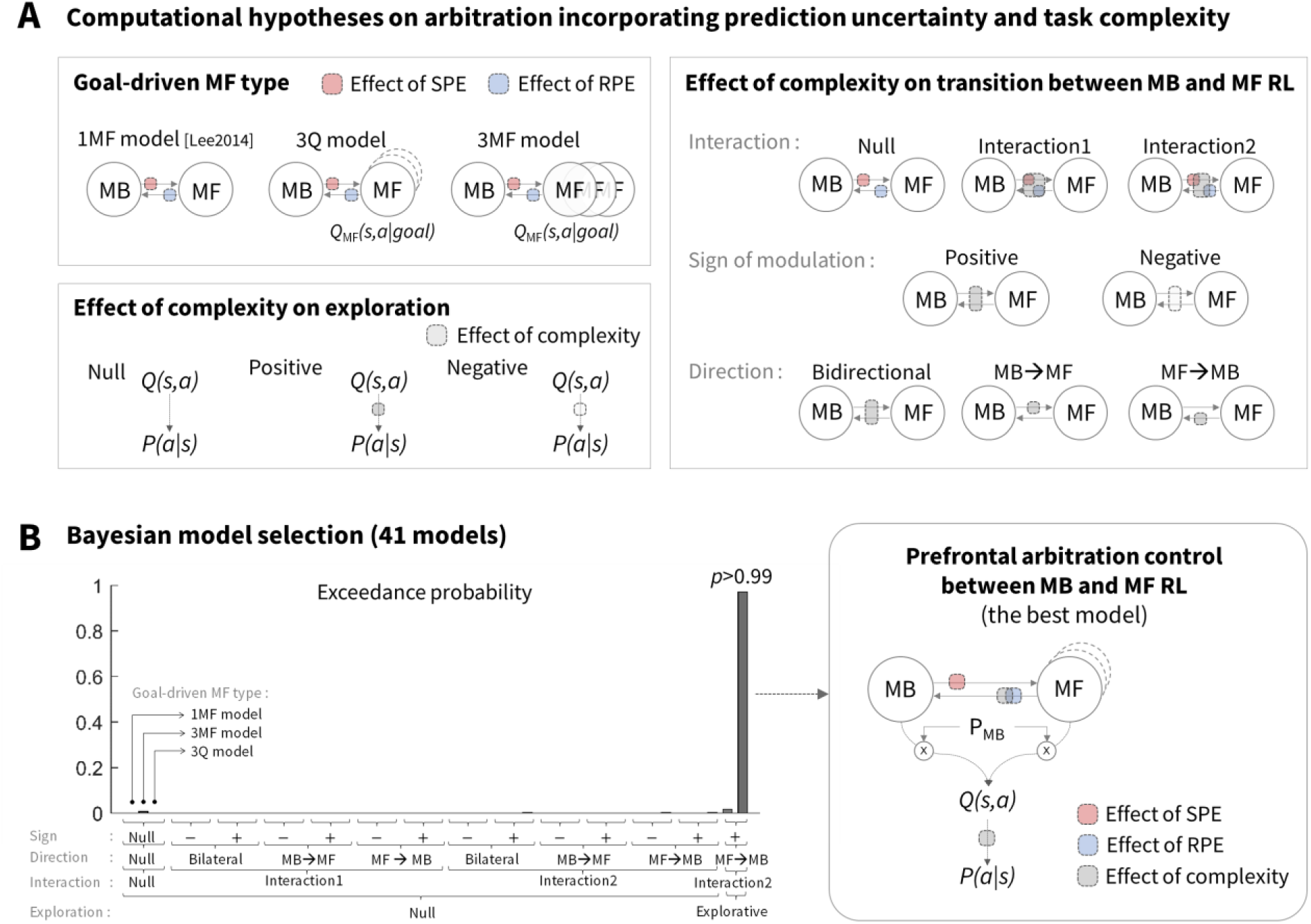
Model comparison analysis on behavioral data. **(A)** We ran a large scale Bayesian model selection analysis to compare different versions of arbitration control. These model variants were broadly classified as reflecting the effect of complexity on the transition between MB and MF RL (13 = 1 + 2×2×3 types), the effect of effect of complexity on exploration (3 variants), and the form of the MF controller (3 variants) each of which is classified by a type of goal-driven MF (3 types), an effect of complexity on transition between MB and MF RL, and an effect of complexity on exploration (3 types). Lee2014 refers to the original arbitration model (Lee et al., 2014). **(B)** Results of the Bayesian model selection analysis. Among a total of 117 versions, we show only 41 major cases for simplicity, including the original arbitration model and other 40 different versions that show non-trivial performance, (but the same result holds if running the full model comparison across the 117 versions). The model that best accounts for behavior is the version {3Q model, interaction type2, excitatory modulation on MF→MB, explorative} (exceedance probability > 0.99; model parameters are shown in the supplementary Table S1).

We found that one specific model variant provided a dominant account of the behavioral data, with an exceedance probability of 0.99. In the best fitting model variant, an increase in task complexity exerted a positive modulatory effect on the transition between model-free and model-based control. That is, the best-fitting model supported an effect of complexity on arbitration such that an increase in complexity produced an *increased* tendency to transition from model-free to model-based control. Recall, that this is not compatible with our initial hypothesis that increased complexity would generally tax the accuracy of the model-based controller, thereby resulting in an increase in model-free control. However, the best-fitting model also proscribed that uncertainty and complexity interact, such that under conditions of both high uncertainty AND high complexity, model-free control would become favored. We will describe in more detail the nature of this interaction in the section below. Secondly, the best-fitting model had the feature that increasing complexity increases the degree of exploration, favoring the hypothesis that subjects tend to explore more under conditions of high-task complexity. Finally, the best fitting model-variant also had the feature that the state-space for the model-free agent was sub-divided according to which goal was currently selected (assuming that the goal selected corresponded to the maximum token value) i.e. the 3Q model, as opposed to a single model-free agent that ignores token values, or separate model-free agents. In summary, we found evidence for an arbitration model that assumes an effect of both uncertainty and complexity on arbitration between MB and MF RL, and furthermore that these two variables interact to drive arbitration as detailed in the following section.

### Tradeoff between model-based and model-free reinforcement learning in the presence of uncertainty and complexity

To gain a better insight into the role of uncertainty and complexity together on choice of the MB vs MF RL strategy according to the best fitting model, we examined the behavior of the model for the four blocked experimental conditions: low/high uncertainty x low/high complexity (see Figure 4A). For this we examined the model weight P_MB_ which is the (dynamically changing) weight assigned to the model-based controller through the arbitration process on a trial by trial basis according to the best fitting model. This P_MB_ weight was binned and averaged within each subject according to whether or not the trial was high or low in complexity and high or low in state-space uncertainty, and the fitted model-weights were then averaged across participants. Note that these are model fits, and thereby illustrate the behavior of the model as fit to the behavioral data, rather than being directly informative about participants’ actual behavior. In essence, this is a way to understand the behavioral predictions of the model itself. When interrogating the fitted model in this way we found an effect of uncertainty and complexity on the weighting between model-based and model-free control (Figure 4A; 2-way repeated measures ANOVA; *p*<1e-4 for the main effect of both state-transition uncertainty and task complexity; *p*=0.039 for the interaction effect). Specifically, according to the model, MB control is preferred when the degree of task complexity increases, whereas MF is favored when the amount of state-space uncertainty increases. A more intriguing finding is that the increase in state-transition uncertainty tends to nullify the effect of task complexity or vice versa.

**Figure 4.**
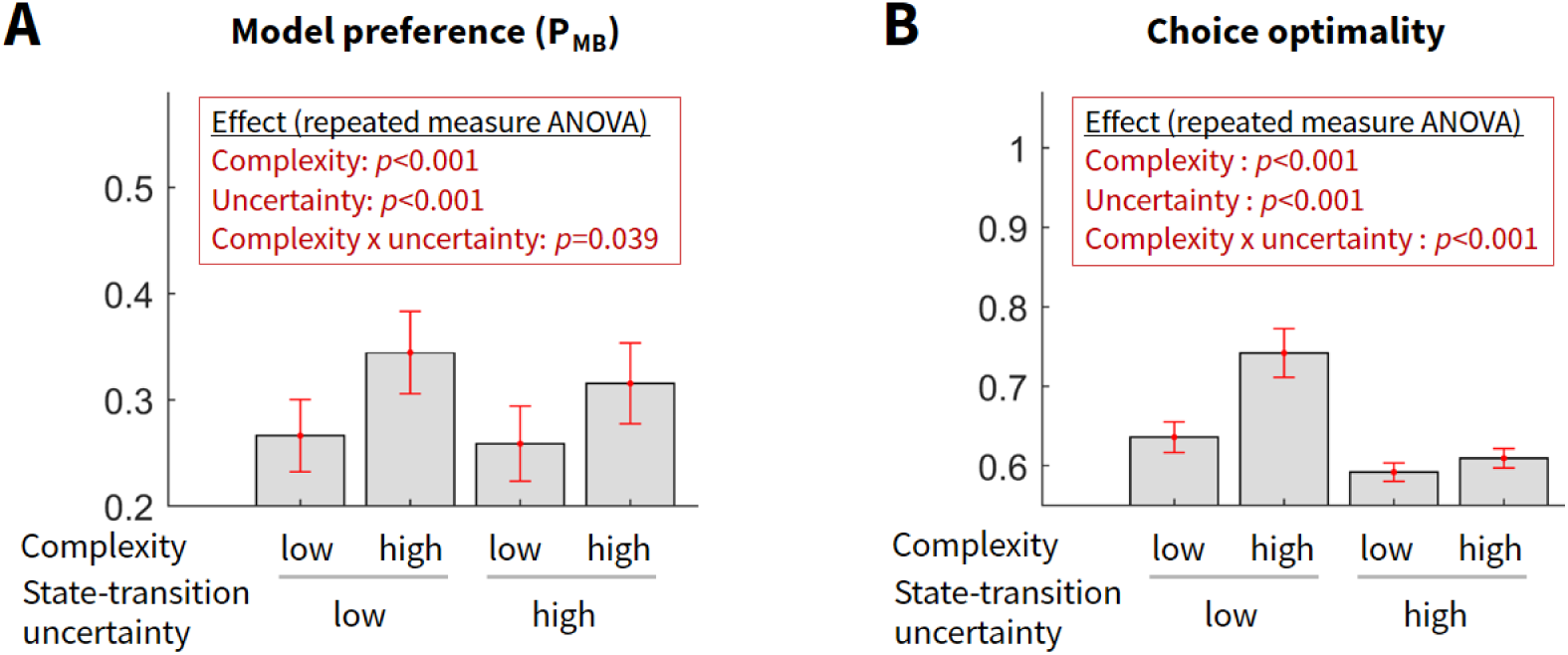
**(A)** Degree of model-based engagement (weighting allocated to the MB strategy; P_MB_) predicted by our computational model of arbitration control (Figure 3B). For this, we ran a deterministic simulation in which our computational model experiences exactly the same episode of events as each individual subject, and we generated the trial-by-trial output: the model-based weight (P_MB_). Error bars are SEM across subjects. **(B)** Participants’ choice optimality. To compute the choice optimality, we computed the degree of match between subjects’ actual choice and an ideal agent’s choice corrected for the number of options. For example, the max value 1 refers to the case in which a subject makes the same choice as the ideal agent’s. The ideal agent is assumed to have a full access to information of state-transition uncertainty and task complexity. Both of the model preference and the choice optimality were calculated for the four experimental conditions (low/high state-transition uncertainty x low/high task complexity). Shown in red boxes are the effect of the two experimental variables on each measure (2-way repeated measures ANOVA). Error bars are SEM across subjects.

To provide a more direct test of the extent to which participants actual behavior is consistent with the arbitration model predictions, we defined an independent behavioral measure which we call choice optimality. Choice optimality quantifies the extent to which participants on a given trial took the objectively best choice had they complete access to the task state-space, and a perfect ability to plan actions in that state-space. It is defined as the ratio of trials on which the subject’s choice matches with the choice of the ideal agent assumed to have a full access to information of state-transition uncertainty and task complexity. Choice optimality is a good proxy of the extent to which participants engage in model-based control, because in principle assuming complete knowledge of the state-space and no cognitive constraints, the model-based agent will always choose more optimally than a model-free agent. Consistent with the model predictions illustrated in Figure 4A, we found a strong effect of uncertainty and complexity on choice optimality in participant’s actual behavior (Figure 4B; 2-way repeated measures ANOVA; *p*<1e-4 for both the main effect and the interaction effect). Consistent with the prediction of the arbitration control model, the effect of complexity on choice optimality diminishes when the state-space uncertainty becomes high.

In summary, these results provide both a computational and behavioral account of how participants regulate the tradeoff between MB and MF RL in the presence of uncertainty and task complexity: they tend to favor use of a MB RL strategy under conditions of high compared to low task complexity, while at the same time they tend to resort to MF RL when the amount of state-space uncertainty increases to the level at which the MB RL strategy can no longer provide reliable predictions. However, these variables interact such that under conditions of both high complexity and high uncertainty, model-free control is favored over and above the effects of each of these two variables alone.

### Neural representations of model-based and model-free RL

To we ran a general linear model analysis (GLM) on the fMRI data in which each variable of the computational model that best fit behavior is regressed against the fMRI data (see Methods for model-specification details). First, we replicated previous findings indicating neural encoding of prediction error signals, SPE and RPE, the two key variables necessary for the update of the value of the MB and MF RL (For full details, see the supplementary Table S2). Consistent with previous findings, we found that SPE signals were encoded in dorsolateral prefrontal cortex (dlPFC) (*p*<0.05 family-wise error (FWE) corrected) (Gläscher et al., 2010; Lee et al., 2014) and RPE signals in the ventral striatum (significantly at *p*<0.05 small volume corrected) (Lee et al., 2014; McClure et al., 2003; O’Doherty et al., 2003). Second, we replicated neural correlates of the chosen value signal for the MB and MF controllers. The MB value signal was associated with BOLD activity in multiple areas within the PFC (*p*<0.05 cluster-level corrected), whereas the MF value signal was found in supplementary motor area (SMA) (*p*<0.05 FWE corrected) and notably posterior putamen (significantly at *p*<0.05 small volume corrected), which has previously been implicated in MF valuation (Tricomi et al., 2009; Wunderlich et al., 2012). Third, we found evidence in the ventromedial prefrontal cortex (vmPFC) for an integrated value signal that combines model-based and model-free value predictions according to their weighted combination as determined by the arbitration process (Q(s,a) shown in Figure 3B; significantly at *p*<0.05 small volume corrected). In addition, we found evidence for the implementation of the goal-driven MF model (Figure S2), which is that the activity of medial frontal gyrus was found to be bilaterally correlated with the goal change signals (*p*<0.05 FWE corrected), the information necessary for the agent to determine whether it caches out a MF value signal in order to achieve a new goal.

### Arbitration signals in prefrontal cortex

We then examined the fMRI data for evidence of signals pertaining to the arbitration process. Replicating our previous results (Lee et al., 2014), we found evidence that the inferior lateral prefrontal cortex (ilPFC) bilaterally encodes reliability signals for both the MB and the MF controllers alone (Fig 5A). But what we found to be best correlated with the activity of ilPFC is the maximum of the reliability of the MB and MF systems, that is, when using a regressor in which the reliability value of whichever system was most reliable on a given trial is input as the value for that trial (Figure 5A; *p*<0.05 cluster-level corrected). These findings are again fully consistent with our previous findings (Lee et al., 2014).

**Figure 5.**
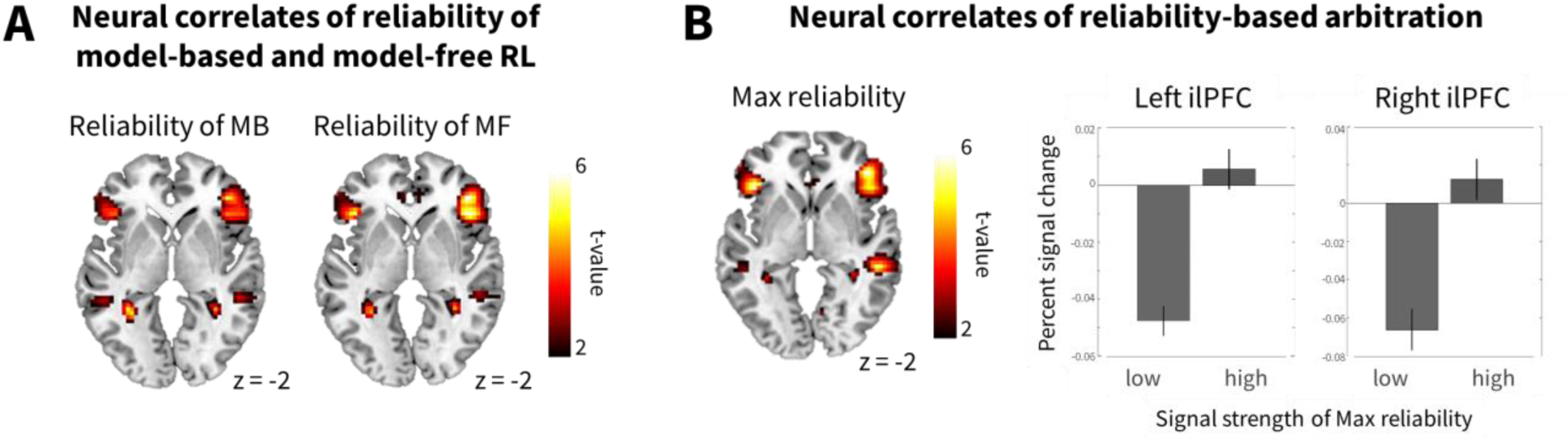
Neural signatures of model-free and model-based reinforcement learning and arbitration control. **(A)** Bilateral ilPFC encodes reliability signals for both the MB and the MF systems. Note that the two signals are not highly correlated (absolute mean correlation < 0.3); this task design was previously shown to successfully dissociate the two types of RL (Lee et al., 2014). Threshold is set at p<0.005. **(B)** (Left) Inferior-lateral prefrontal cortex bilaterally encodes reliability information on each trial of both MB and MF RL, as well as whichever strategy that provides more accurate predictions (“max reliability” (Lee et al., 2014)). (Right) The mean percent signal change for a parametric modulator encoding a max reliability signal in the inferior lateral prefrontal cortex (lPFC). The signal has been split into two equal-sized bins according to the 50th and 100th percentile. The error bars are SEM across subjects.

### Model comparison against fMRI data

Next we aimed to formally test our hypothesis of uncertainty and complexity-sensitive arbitration control against the previous hypothesis that arbitration control is based solely on changes in prediction uncertainty alone (Lee et al., 2014). For this we compared two separate arbitration models against the fMRI data. One was the best fitting model described above in which both task complexity and reliability are taken into account as playing a role in driving the arbitration process. The second, was a model in which only reliability was involved in the arbitration process, as first utilized by Lee et al. (Lee et al., 2014). We then compared the fit of these two models to the fMRI data in two brain regions, ilPFC for the reliability signals and vmPFC for the valuation signals. For this we ran a Bayesian model selection analysis (Stephan et al., 2009), using spherical ROIs centered on the coordinates from our 2014 study, thereby ensuring independence of the ROI selection from the current dataset. In a majority of voxels in both regions of interest, reliability signals from the model incorporating both reliability and task complexity were preferred over the previous model incorporating reliability only (Figure 6). These findings support our hypothesis that the model in which complexity is taken into account provides a better account of prefrontal mediated arbitration control.

**Figure 6.**
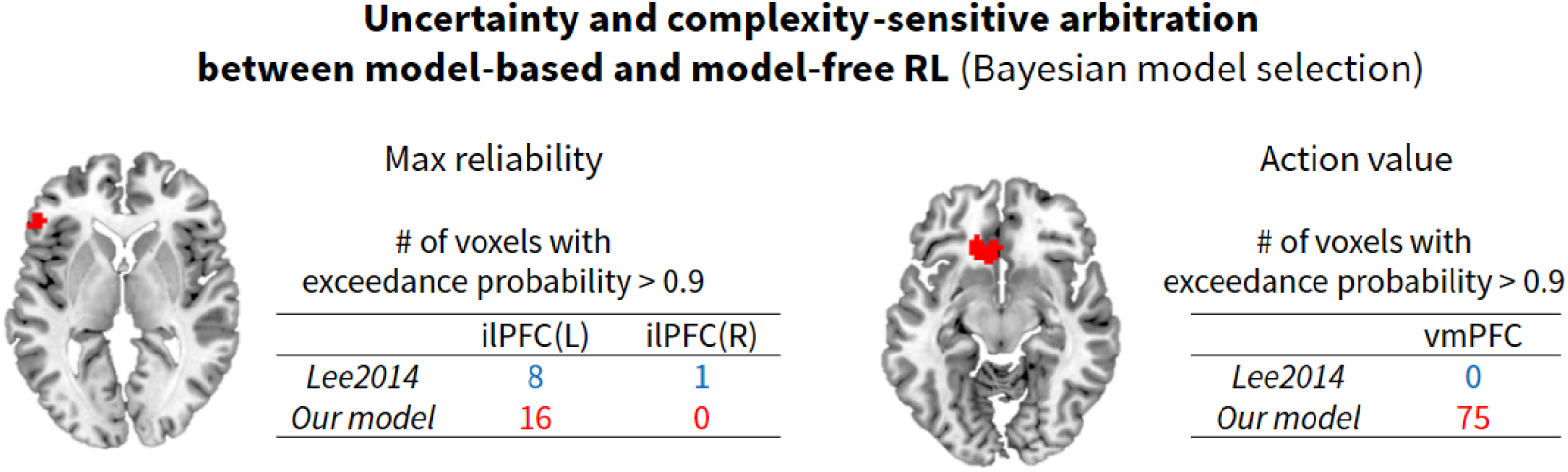
Results of a Bayesian model selection analysis. The red blobs and table show the voxels and the number of voxels, respectively, that favor each model with an exceedance probability > 0.95, indicating that the corresponding model provides a significantly better account for the BOLD activity in that region. *Lee2014* refers to an arbitration control that takes into account only uncertainty as used by (Lee et al., 2014), *Current model* refers to the arbitration control model that was selected in the model comparison based on the behavioral data which incorporates both prediction uncertainty and task complexity. For an unbiased test, the coordinates of the ilPFC and the vmPFC ROIs were taken from (Lee et al., 2014).

### Modulation of inferior prefrontal reliability signal by complexity

The above findings demonstrate that reliability signals from an arbitration model that takes into account task complexity, provides a better account of fMRI activity than reliability signals derived from a model that does not incorporate complexity, reflecting an implicit contribution of complexity to the arbitration process. However, these findings do not provide direct evidence for an explicit representation of task complexity in the ventrolateral prefrontal arbitration region. To test for an explicit contribution of task complexity to the arbitration signal, we ran an additional fMRI analysis in which we included a parametric regressor denoting the onset of trials of high task vs low task complexity as a separate regressor of interest. We then entered another additional regressor, which corresponds to the formal interaction of Max reliability with task complexity. That is, to test for areas in which the reliability signal was modulated differently depending on whether a specific trial was high or low in complexity. While we found no significant effect of the main effect of task complexity in our main regions of interest (Table S2), we found evidence for a significant interaction effect of complexity and reliability. Specifically, a region of inferior lateral prefrontal cortex bilaterally was found to show a significant negative interaction between complexity and reliability (Fig. 7A). This region was found to overlap substantively with the regions of inferior lateral prefrontal cortex found to exhibit a main effect of reliability (Fig. 7B). To visualize the effect of the interaction in this region we in a post-hoc analysis, extracted the average % signal change from the clusters exhibiting the interaction in left and right ilPFC respectively. We binned the signal according to whether reliability was high or low, and whether complexity was high or low, shown in Fig. 7B. Reliability signals were plotted separately for model-free and model-based reliability, although the results are similar for max reliability. As can be seen, the reliability signals show evidence of being attenuated particularly when complexity is high relative to when complexity is low. This provides evidence that these two signals relevant for driving arbitration interact with each other in ilPFC.

**Figure 7.**
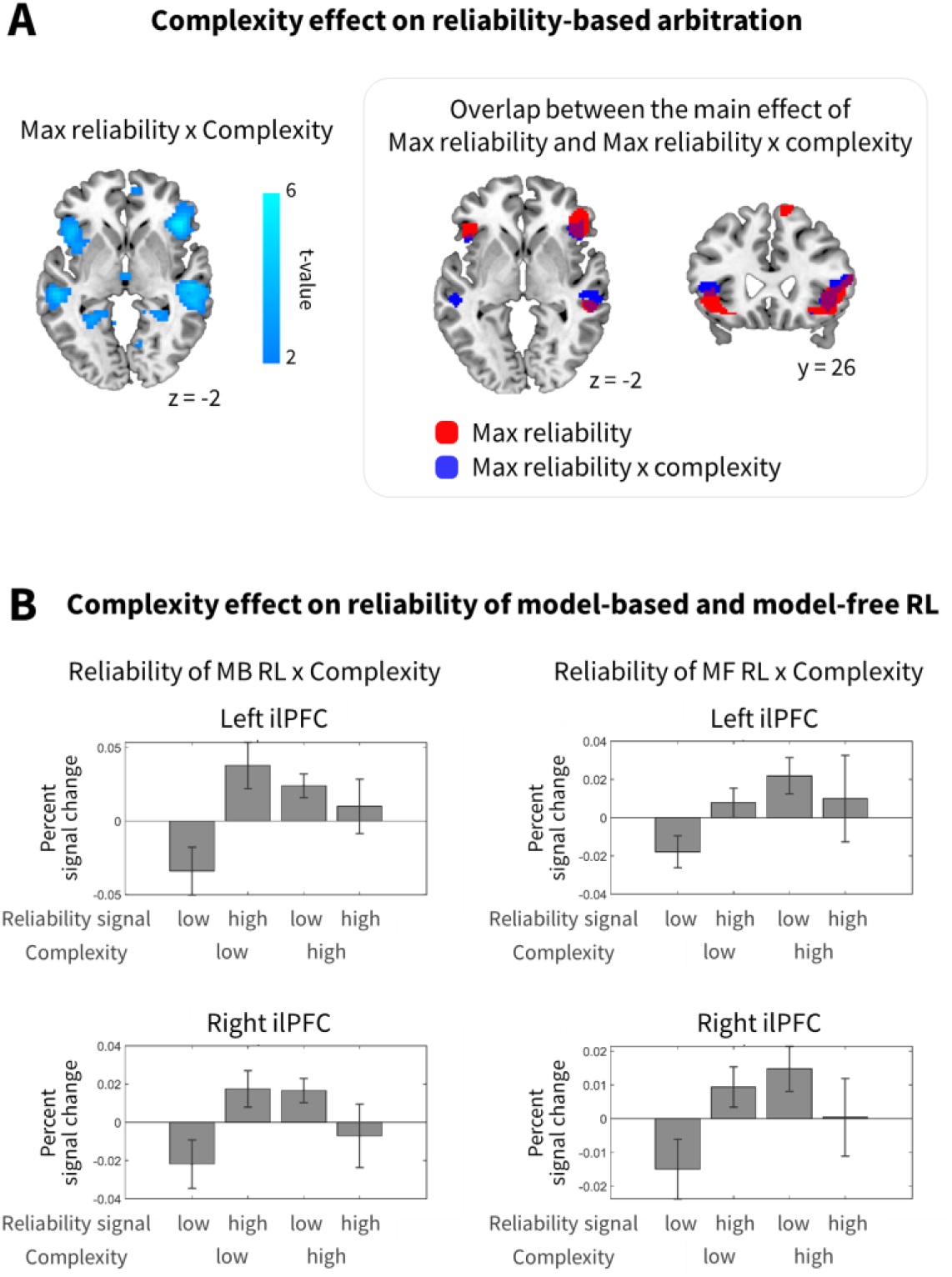
Bilateral ilPFC was found to exhibit a significant interaction between complexity and reliability (Max reliability x complexity). Statistical significance of the negative effects is illustrated by the cyan colormap. The threshold is set at p<0.005. (Right) The brain region reflecting the interaction effect largely overlaps with the brain area implicated in arbitration control. The red and blue regions refers to the main effect of max reliability and the interaction between reliability and task complexity, respectively, thresholded at p<0.001. **(B)** Plot of average signal change extracted from left and right ilPFC clusters showing the interaction, shown separately for reliability signals derived from the MF and MB controllers. Data are split into two equal-sized bins according to the 50th and 100th percentile of the reliability signal, and shown for the trials in the low and high complexity condition separately. The error bars are SEM across subjects.

## DISCUSSION

We provide evidence supporting the interaction of two key variables in driving arbitration of control between model-based and model-free reinforcement learning. In addition to replicating our previous finding implicating the uncertainty (or reliability) of the predictions made by the model-based and model-free controllers in moderating the influence of these two systems over behavior, we found evidence that the complexity of the state-space, which is a contributor to the computational cost faced by the model-based controller, also contributes to the setting of the balance between these two systems. These behavioral findings were supported by evidence that a region of the brain previously implicated in the arbitration process, the inferior lateral prefrontal cortex not only encodes signals related to the reliability of the predictions of the two systems that would support an uncertainty-based arbitration mechanism, but furthermore that activity in this region is better accounted for by an arbitration model that also incorporates the effects of task complexity into the arbitration process. Moreover, we found evidence that task complexity and reliability appear to directly interact in this region. Taken together, these findings help to advance our understanding of the contribution of two key variables to the arbitration process at behavioral and neural levels.

We found direct evidence for a contribution of task complexity on arbitration. First, in our large scale model comparison we found empirical support for a version of the arbitration process in which the complexity variable has a positive modulation effect on the transitions from MF to MB. Second, this is corroborated by the fact that the best fitting model exhibited an increased preference for MB over MF in the high complexity condition on average. Third, in an independent behavioral analysis which uses choice optimality to quantify the extent to which choice behavior is guided by MB RL, we found that subjects’ choice optimality increases with the degree of task complexity. These results together suggest that an increase in task complexity creates an overall bias towards MB RL, contrary to our initial hypothesis in which we considered that increased complexity would tax the model-based system resulting in increased model-free control. Another interesting finding supporting this idea is that an increase in task complexity makes choices more flexible and explorative. In summary, these findings suggest that humans attempt to resolve task complexity by engaging a more explorative MB RL strategy.

We also found an effect of state-space uncertainty on the arbitration process. Specifically, very high state-space uncertainty makes subjects resort more to a MF RL strategy. This effect arises because high state-space uncertainty results in a lowered reliability of the predictions of the model-based controller, thereby resulting in a reduced contribution of behavior of the model-based controller. It should be noted that, the model-based controller should generally compute a more accurate prediction than its model-free counterpart, which by contrast necessarily generates approximate value predictions. However, this holds only under the situation where the model-based controller has access to a reliable model of the state-space. If its state-space model is not reliable or accurate, then the model-based controller cannot generate accurate predictions about the value of different actions. In this task, we influence the extent to which the model-based controller has access to reliable predictions about the state-space by directly modulating the state-space uncertainty. Thus, under conditions in which the state-space model is highly unreliable, humans appear to rely more on model-free control. Conversely, as we have shown previously (Lee et al., 2014), if the reliability of the model-free controller is decreased, participants will all else being equal rely more on model-based control. Thus, the uncertainty in the predictions of these controllers, appears to play a key role in underpinning the arbitration process between them.

In addition to the main effects of complexity and state-space uncertainty, we have shown that these two variables interact. Under conditions where both state-space uncertainty is high and complexity is high, the model-based system appears to be disproportionately affected, in that participants abandon model-based control in favor of model-free control. Thus, our hypothesis about an effect of complexity resulting in decreased model-based control was borne out in a qualified manner, in that this effect only happens when state-space uncertainty is high. This finding suggests that the arbitration process takes into account the effects of both of these variables at the same time, and dynamically finds a tradeoff between them. Participants appear to use the MB RL strategy to resolve uncertainty and complexity, but owing to the fact that MB RL is more cognitively demanding than MF RL, they resort to the default strategy, MF RL, when the performance gain of MB RL does not outweigh the level of cognitive load required for MB RL.

The present findings are also relevant to the predictions of expected value of control (EVC) theory (Narayanan et al., 2013; Shenhav et al., 2013). One prediction of the EVC theory is that increased control signal intensity would lock the MB system in, thereby letting the MF system take over control over behavior. Another related finding is that MB learning with prior training remains intact under cognitive load (Economides et al., 2015). While the theory predicts that increasing task difficulty brings about increase in the intensity of the cognitive control signal, the theory itself does not offer any predictions about how control signal intensity influences RL. Our computational model explains how the brain chooses between MB and MF RL with varying amount of cognitive control intensity, and furthermore, why this choice is made.

The model comparison analysis of the present study also revealed that task complexity affects transitions from MF to MB RL, but not transitions in the other direction. This finding provides further evidence to support the existence of an asymmetry in arbitration control such that arbitration is performed in a way that selectively gates the MF system (Lee et al., 2014; Wunderlich et al., 2012). These results may be reasonable from an evolutionary perspective in that the implementation of MF learning in parts of the basal ganglia may have arisen earlier on in the evolutionary history of adaptive intelligence, while later on, cortically mediated MB control may have emerged so as to deal with more complex situations.

Our study also advances understanding into the nature of the computations being implemented in the inferior prefrontal cortex during the arbitration process. This region was found to correlate not only with the reliability of the predictions of the two control systems, as shown previously (Lee et al., 2014), but this region also was found to incorporate information about task complexity. The reliability signal itself was found to reflect the effects of task complexity, as shown by the formal model comparison in which activity in this region was better accounted for by an arbitration model that incorporated task complexity compared to a model that ignores task complexity. Moreover, we found that when testing directly for an interaction between the reliability signals and complexity, an overlapping region of inferior prefrontal cortex was found to show evidence for a significant interaction between these signals. These findings therefore demonstrate that task complexity directly modulates the putative neural correlates of the arbitration process.

While our findings advance our understanding of the nature of the arbitration process, it is also important to acknowledge that a number of open questions remain. A fundamental open question concerns how the arbitration computations within inferior prefrontal cortex are actually implemented at the neuronal level. While our findings show correlations between various arbitration-related signals in this region such as reliability and complexity, how these signals are utilized at the neural level to implement the arbitration process is not yet addressed. Building on the present study and earlier studies investigating executive control mechanisms in prefrontal-striatal circuitry (Aron et al., 2003, 2004; Balleine and O’Doherty, 2010; Burguière et al., 2013; Cockburn and Frank, 2011; Coutureau and Killcross, 2003; Daw et al., 2011; Donoso et al., 2014; Gremel and Costa, 2013; Koechlin et al., 2003; Robbins, 1996; Rushworth et al., 2011; Smith et al., 2012; Tanji and Hoshi, 2008), more biologically plausible models of the arbitration process will need to be developed to go beyond the algorithmic level used in the present study. Furthermore, in order to guide the development of such models it will ultimately be important to get a better understanding of the underlying neuronal dynamics in these prefrontal regions during the arbitration process, using techniques with better spatial and temporal resolution than available with fMRI.

An important limitation of the task we have used here, is that we have studied behavior under conditions of high instability and/or variability in terms of the transitions between model-based and model-free control. This is done by design, because to detect arbitration related signals both behaviorally and neurally within the framework of an fMRI study, we needed to maximize the variance in the transition between these two different forms of behavioral control. However, in real-world behavior, it would be expected that the transitions between model-based and model-free control would typically be evolving at a slower pace. One of the main advantages of model-free control is the lower computational cost entailed by engaging cached values learned via model-free RL compared to model-based RL. A natural consequence of this is that the model-based system should cease computing action-values when the model-free system is in control, as otherwise the computational cost advantage gained by increasing model-free control would be moot. A limitation of the present arbitration model is that it assumes that model-based values continue to be computed throughout the task. This is so because we did not find behavioral evidence that such signals ceased to be estimated during the task, which would be manifested at the behavioral level by a complete dominance of model-free control over behavior. However, we suspect that in real-world behavior, the model-based system would eventually go offline, and this should be reflected ultimately in behavioral dominance of model-free control. More generally, it will be important to study the behavior of model-based and model-free controllers across a wide range of tasks and experimental conditions in order to gain a more complete understanding of the nature of the arbitration process.

Finally, it should be noted that the cognitive complexity manipulation we used here by which we increased the number of available state-spaces available in the decision problem, can also impact on the model-free system, because the increase in the number of actions that need to be learned, means that the model-free system has less opportunity to sample those state-action pairs thereby having less opportunity to acquire accurate value representations. Thus, the trade-off faced by the model-based and model-free system under these conditions is more complicated than merely reflecting the sole effects of computational cost on the model-based system. Future studies could therefore focus on more clearly separating the effects of computational load from sampling history.

In conclusion, we provide behavioral and neural evidence for the effect of prediction uncertainty and task complexity on RL. First, the present findings provide a theoretical insight into how the brain dynamically combines different RL strategies to deal with uncertainty and complexity. Second, the findings help us decipher the asymmetrical nature of arbitration control – supporting the notion that the primary function of arbitration control is to regulate the MF system, as opposed to exerting control on both systems to an equal extent. Third, we found that such an arbitration control principle is best reflected in neural activity patterns in the inferior lateral prefrontal cortex, the same area we previously found to play a pivotal role in arbitration control, thereby significantly advancing our understanding of the theory of prefrontal arbitration between MB and MF RL. Taken together, the findings in the present study foster a deeper appreciation of the role of ventrolateral prefrontal cortex in arbitration control.

### Experimental Procedures

#### Participants

Twenty four right-handed volunteers (ten females, with an age range between 19 and 55) participated in the study, 22 of whom were scanned with fMRI. They were screened prior to the experiment to exclude those with a history of neurological or psychiatric illness. All participants gave informed consent, and the study was approved by the Institutional Review Board of the California Institute of Technology.

#### Stimuli

The image set for the experiment consisted of 126 fractal images to represent states, three kinds of color coins (red, blue, and silver), and four kinds of fractal images to represent outcome states associated with each color coin (red, blue, silver, and none). The colors of the outcome state image were accompanied by numerical amounts which indicate the amount of money that subjects could receive in that state. Before the experiment began, the stimulus computer randomly chose a subset of eleven fractal images to be subsequently used to represent each state in that specific participant.

#### Task

Participants performed a sequential two-choice Markov decision task, in which they need to make two sequential choices (by pressing one of four buttons: L1, L2, R1, R2) to obtain a monetary outcome (token) at the end stage. Making no choice in 4 seconds had a computer make a random choice to proceed and that trial was marked as a penalizing trial. Each trial begins with a presentation of values of each token in that trial, followed by a presentation a fractal image representing a starting state. The presentation of each state is accompanied by choice availability information shown at the bottom of the screen. Only two choices (L1 and R1) are available in the starting state (S1). The starting state is the same across all trials. Making a choice in the starting state is followed by a presentation of another fractal image representing one of ten states (S2-S11). The states were intersected by a variable temporal interval drawn from a uniform distribution between 1 to 4 seconds. The inter-trial interval was also sampled from a uniform distribution between 1 to 4 seconds. The reward was displayed for 2 seconds. At the beginning of the experiment, subjects were informed that they need to learn about the states and corresponding outcomes to collect as many coins as possible and that they will get to keep the money they cumulatively earned at the end of the experiment. Participants were not informed about the specific state-transition probabilities used in the task except they were told that the contingencies might change during the course of the experiment. In the pre-training session, to give participants an opportunity to learn about the task structure, they were given 100 trials in which they can freely navigate the state space by making any choices. During this session, the state-transition probability was fixed at (0.5,0.5) and the values of all color tokens are fixed at 5, indicating that any token color would yield the same amount of monetary reward. The experiment proceeded in five separate scanning sessions of 80 trials each on average.

In order to effectively dissociate the model-free strategy from the model-based, the experimental design of the present study introduces two task parameters: specific goal-condition and state transition probabilities. First, to create a situation in which the model-based control is preferred over the model-free control, the present experimental design introduced a generalized version of the specific goal-condition (Lee et al., 2014), in which all token values are randomly drawn from a uniform probability distribution U(1,10) from trial-to-trial. If participants reached the outcome state associated with a token, then they would gain the corresponding monetary amount. Note that this goal-value manipulation is intended to encourage participants to act on a stable model-based control strategy, as opposed to developing separate multiple model-free strategies for each color tokens in the absence of the model-based control.

Second, to create a situation where the model-free control overrides the model-based control and to further dissociate the model-free from the model-based, changes to the state transition probabilities were implemented. Two types of state-transition probability were used – (0.9,0.1) and (0.5,0.5) (a *low* and a *high state-transition uncertainty condition*, respectively). They are the probabilities that the choice is followed by going into the two consecutive states. For example, if you make a left choice at the state 1 and the state transition probability is (0.9,0.1) at that moment, then the probability of your next state being state 2 is 0.9 and the probability for state 3 is 0.1. The order of the block conditions was randomized. The blocks with the state transition probability (0.9,0.1) consists of three to five trials, whereas those with (0.5,0.5) consists of five to seven trials; it was previously shown that with (0.9,0.1) participants feel that the state transition is congruent with the choice, whereas with (0.5,0.5) the state transition is random (Lee et al., 2014). Furthermore, the changes at these rates ensures that tonically varying changes in model-based vs model-free control can be detected at experimental frequencies appropriate for fMRI data. The state-transition probability value was not informed to participants; estimation of state-transition probabilities can be made by using the model-based strategy.

To manipulate the state-space complexity, the present study also introduced the third task parameter, the number of available choices. Two types of choice sets were used – (L, R) and (L1, L2, R1, R2) (a *low* and a *high state-space complexity condition*, respectively). The order of the block conditions was randomized. To preclude the task being too complex, changes in the number of available choices occur only in the second stage of each trial, while in the first stage the number is always limited to two (L and R).

#### Computational model of arbitration

Computational models of arbitration used in this study are based on the previous proposal of arbitration control (Lee et al., 2014). The original arbitration model uses a dynamic two-state transition (Dayan and Abbott, 2001) to determine the extent to which the control is allocated to a model-based learner (MB) (Gläscher et al., 2010) and to a model-free SARSA learner (MF) (Sutton and Barto, 1998) at each moment in time. Specifically, the change of the control weight *P*_MB_ (the probability of choosing a model-based strategy) is given by the difference between two types of transition: MF→MB and MB→MF:

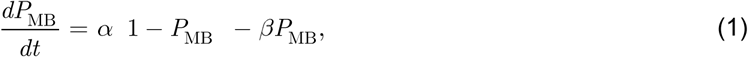

Where *α*, *β* refers to the transition rate MF→MB and MB→MF, respectively.

The transition rate *α* (MF→MB) is found to be a function of reliability of the MF strategy that reflects the average amount of reward prediction error (Lee et al., 2014):

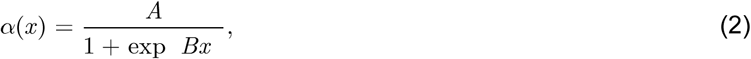

where *x* refers to MF reliability and the two free parameters *A*, *B* refers to the maximum transition rate and the steepness, respectively. Likewise, the transition rate *β* (MB→MF) is defined as a function of MB reliability that reflects the posterior estimation of the amount of state prediction error (Lee et al., 2014).

### Computational hypotheses on arbitration incorporating prediction uncertainty and task complexity

#### 1. Goal-driven MF type

To test the hypothesis on ‘goal-driven MF’ (Figure 3A), we implemented the following versions.

1. 1MF model : refers to the null hypothesis that the MB and the goal-independent MF interacts.
2. 3Q model : refers to the hypothesis that the MB interacts with a single MF with goal-dependent state-action value sets. Specifically, the MF learns a state x action x goal(red/blue/silver) value matrix with a single learning rate.
3. 3MF model : refers to the hypothesis that the MB interacts with goal-dependent multiple MFs. Specifically, each goal is associated with an independent MF (red, blue, and silver) with a separate learning rate.

#### 2. Effect of complexity on transition between MB and MF RL

To test the effect of the state-space complexity on arbitration control, we define a transition rate as a function of both reliability and complexity (see the right box of Figure 3A). The following variants of the transition function (2) were used.

1. Type of interaction For simplicity, we only show the variants of the transition rate *α* (MF→MB). The same rule can be applicable to the transition rate *β*. -Null : assumes that there is no complexity effect on arbitration control; refer to the equation (2). -Interaction1 : assumes that there is a direct interaction effect (complexity (z) x reliability (x)) on arbitration control.

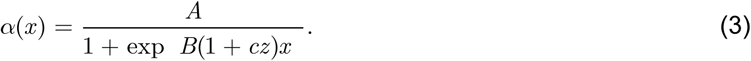 -Interaction2 : assumes that there is a indirect interaction effect on arbitration control. Although there is no interaction term (zx), the transition rate is a function of both complexity (z) and reliability (x).

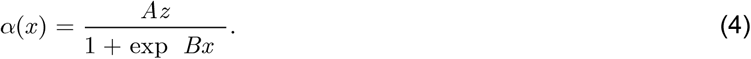
2. Sign of modulation -Positive : assumes that the complexity has a positive influence on the transition rate. We set z=1 and 2 for a low and high complexity condition, respectively. -Negative : assumes that the complexity has a negative influence on the transition rate. We set z=2 and 1 for a low and high complexity condition, respectively.
3. Direction of modulation Bidirectional : assumes that the complexity influence the both transition rates MB→MF and MF→MB. This means that the above rules (the type of interaction and the sign of modulation) are applied to both transition rates. -MB→MF : assumes that the complexity influence the transition rates MB→MF only. -MF→MB : assumes that the complexity influence the both transition rates MF→MB only.

To test the effect of the state-space complexity on exploration, we define an exploration parameter as a function of complexity (see the bottom-left box of Figure 3A).

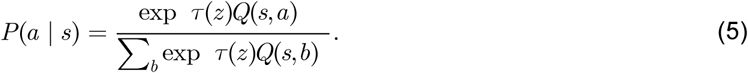

#### 3. Effect of complexity on exploration

To test the hypothesis that increasing complexity increases the degree of explorative choices (Figure 3A), we set τ(*z*) = 1 *and* 0.5 for the low and high complexity condition, respectively. For testing the hypothesis that increasing complexity decreases the degree of explorative choices, we set τ(*z*) = 0.5 *and* 1.

Note that we compared prediction performance of all combinations of the above cases, and for simplicity we showed the results of only 41 major cases (Figure 3B); in most of cases prediction performance is far below than the stringent threshold (exceedance probability *p*=1e-3).

#### Relationship between our computational model and a simple mixture of MB and MF RL

In a stable environment (i.e., a fixed amount of state-transition uncertainty and a fixed level of task complexity), the state of our computational model converges to a fixed point. This is specified by the steady-state model choice probability:

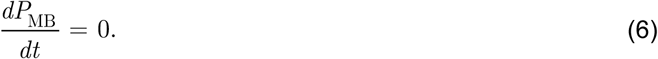

Then by using (1), we get

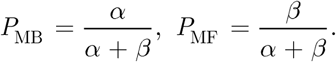

Note that this is the equivalent of a simple mixture of MB and MF RL (Daw et al., 2011).

#### fMRI data acquisition

Functional imaging was performed on a 3T Siemens (Erlangen, Germany) Tim Trio scanner located at the Caltech Brain Imaging Center (Pasadena, CA) with a 32 channel radio frequency coil for all the MR scanning sessions. To reduce the possibility of head movement related artifact, participants’ heads were securely positioned with foam position pillows. High resolution structural images were collected using a standard MPRAGE pulse sequence, providing full brain coverage at a resolution of 1 mm × 1 mm × 1 mm. Functional images were collected at an angle of 30° from the anterior commissure-posterior commissure (AC-PC) axis, which reduced signal dropout in the orbitofrontal cortex. Forty-five slices were acquired at a resolution of 3 mm × 3 mm × 3 mm, providing whole-brain coverage. A one-shot echo-planar imaging (EPI) pulse sequence was used (TR = 2800 ms, TE = 30 ms, FOV = 100 mm, flip angle = 80°).

#### fMRI data analysis

The SPM12 software package was used to analyze the fMRI data (Wellcome Department of Imaging Neuroscience, Institute of Neurology, London, UK). The first four volumes of images were discarded to avoid T1 equilibrium effects. Slice-timing correction was applied to the functional images to adjust for the fact that different slices within each image were acquired at slightly different points in time. Images were corrected for participant motion, spatially transformed to match a standard echo-planar imaging template brain, and smoothed using a 3D Gaussian kernel (6 mm FWHM) to account for anatomical differences between participants. This set of data was then analyzed statistically. A high-pass filter with a cutoff at 129 seconds was used. Full details of the GLM design matrix are provided in Supplementary Methods.

#### Bayesian model selection analyses on fMRI data

To formally test which version of arbitration control provides the best account of responses in inferior lateral prefrontal cortex, we ran a Bayesian model selection (Stephan et al., 2009). We chose three models -*α*, *β*, *α*, *β*_2_*α*_2_, *β* the original arbitration model (Lee et al., 2014) and the two other versions that we found in Bayesian model selection analysis on behavioral data exhibit the second best and the best performance, respectively. We used a spherical ROI centered on the coordinates from the previous study (Lee et al., 2014) with a radius of 10mm.

## Acknowledgements

We thank Peter Dayan for insightful comments and Ralph Lee for his assistance. This work was funded by grants R01DA033077 and R01DA040011 to J.P.O.D. from the National Institute on Drug Abuse. This work was also supported by an Institute for Information & Communications Technology Promotion (IITP) grant funded by the Korea government (No. 2017-0-00451), the ICT R&D program of MSIP/IITP. [2016-0-00563, Research on Adaptive Machine Learning Technology Development for Intelligent Autonomous Digital Companion], the Brain Research Program through the National Research Foundation of Korea(NRF) funded by the Ministry of Science, ICT & Future Planning (NRF-2016M3C7A1914448), the National Research Foundation of Korea(NRF) grant funded by the Korea government(MSIT) (No. 2017R1C1B2008972), the research fund of the Korea Advanced Institute of Science and Technology (KAIST) under Grant code G04150045, and Samsung Research Funding Center of Samsung Electronics under Project Number SRFC-TC1603-06.

## Author contributions

SL and JOD conceived and designed the study. SL implemented the behavioral task and ran the fMRI study. DK, GP, and SL designed computational models and analyzed the data. SL, JOD, and DK wrote the paper. All authors approved the final version for submission.

